# Composition-controlled artificial collagen shows opposing roles of collagen-binding integrins and discoidin domain receptors in neuronal differentiation of PC12 cells

**DOI:** 10.64898/2026.08.25.746976

**Authors:** Kazunori K. Fujii, Kota Tsusaka, Takaki Koide

## Abstract

Collagen, a major component of the extracellular matrix, regulates cellular behaviors, such as adhesion, differentiation, and angiogenesis. These functions are mediated by interactions between specific amino acid motifs within the collagen triple-helical structure and collagen-binding biomolecules. These include cell-surface receptors, such as integrins, discoidin domain receptors (DDRs), and syndecans, a family of transmembrane heparan sulfate proteoglycans (HSPGs). Signals mediated by these receptors are integrated to regulate cell fate. However, native collagen simultaneously presents multiple receptor-binding motifs, making it difficult to isolate receptor-specific functions and to evaluate receptor crosstalk. Here, we introduce a composition-controlled artificial collagen matrix platform that enables independent tuning of multiple receptor-binding motifs within a constant triple-helical scaffold. This material was produced by disulfide crosslinking of chemically synthesized collagen-like triple-helical peptides, each bearing a single defined receptor-binding sequence. By varying the mixing ratios of these peptides before crosslinking, we systematically controlled the composition of receptor-binding motifs within the matrices. We applied this platform to nerve growth factor-dependent neuronal differentiation of PC12 cells, a process supported by collagen. Matrices containing only integrin-binding sequences were sufficient to support this differentiation. Incorporation of an HSPG-binding sequence had little additional effect, whereas incorporation of a DDR-binding sequence suppressed integrin-mediated differentiation and coincided with DDR phosphorylation. These results reveal opposing roles of collagen-binding integrins and DDRs in regulating PC12 cell differentiation. Composition-controlled artificial collagen provides a versatile matrix platform for dissecting functional crosstalk among collagen receptors.

**Graphical abstract:** 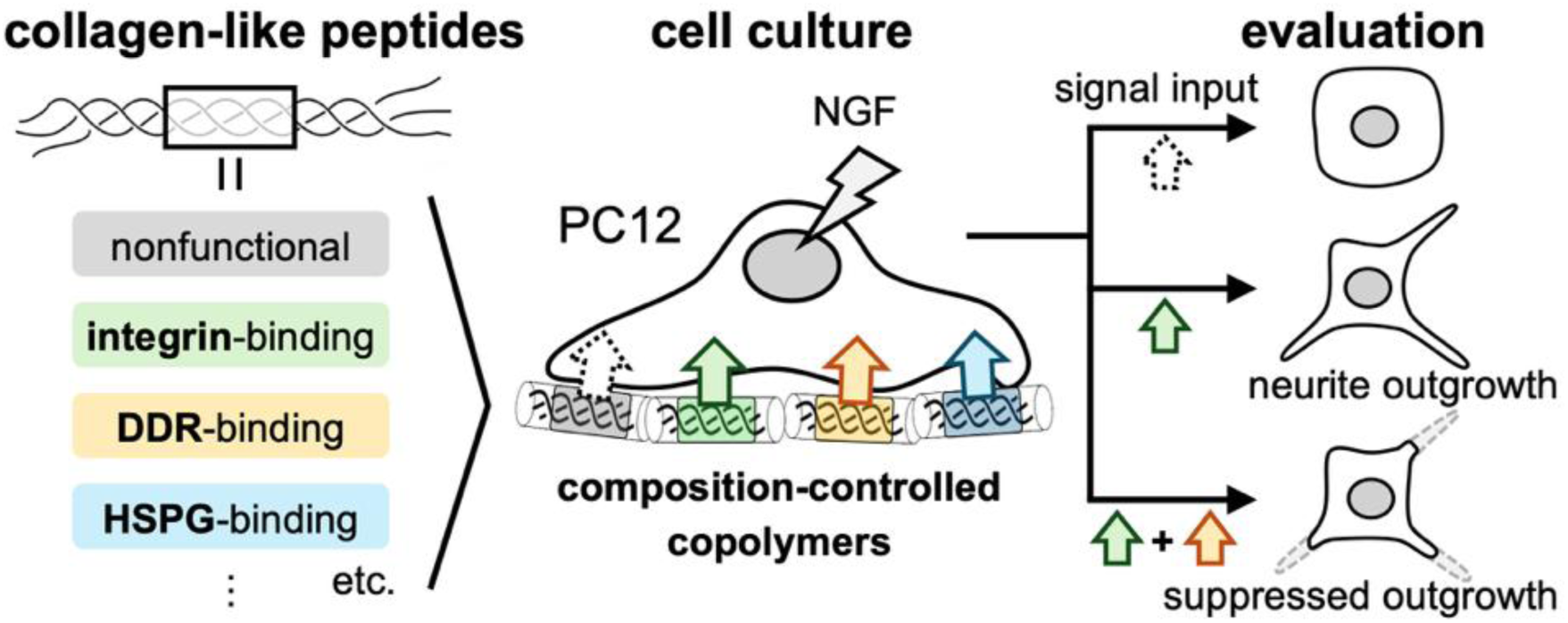

**Statement of significance:** Native collagen presents multiple receptor-binding motifs simultaneously, making it difficult to isolate receptor-specific functions and interrogate receptor crosstalk. We address this limitation with composition-controlled artificial collagen matrices assembled from chemically synthesized collagen-like triple-helical peptides. This approach enables independent, tunable presentation of defined receptor-binding sequences within a constant triple-helical scaffold. Using nerve growth factor-treated PC12 cells as a model, an integrin-binding sequence was sufficient to induce neuronal differentiation. Notably, co-presenting a discoidin domain receptor (DDR)-binding sequence suppressed this response and coincided with DDR activation. This modular matrix strategy enables controlled interrogation of collagen receptor crosstalk and supports the design of cell-instructive biomaterials.

## 1. Introduction

Collagen is a major component of the extracellular matrix and is characterized by a triple-helical structure composed of three polypeptide chains with a repeating Gly–Xaa–Yaa sequence, where Xaa and Yaa can be any amino acid. Proline (Pro) and 4-hydroxyproline (Hyp, O) are frequently found at the Xaa and Yaa positions, respectively. To date, 28 types of collagen have been identified in the human body. Of these, type I collagen is the most abundant. It is a fibrillar collagen with a triple-helical region consisting of 338 repeated Gly–Xaa–Yaa motifs and is often used in collagen research because of its ubiquitous presence in connective tissues. Type IV collagen, in contrast, is a network-forming collagen and a major component of basement membranes.

Collagen interacts with several cell-surface receptors, including collagen-binding integrins (integrins α1β1, α2β1, α10β1, and α11β1) [1], discoidin domain receptors (DDR1 and DDR2), a family of receptor tyrosine kinases [2], and syndecans, a family of transmembrane heparan sulfate proteoglycans (HSPGs) [3]. These interactions are mediated, at least in part, by specific amino acid sequences presented within the triple helix and regulate biological processes such as cell adhesion, differentiation, and angiogenesis. When multiple collagen receptors are engaged, crosstalk among these receptors may regulate cellular processes such as proliferation and differentiation [4]. For instance, DDR1 overexpression in Madin–Darby canine kidney cells suppresses integrin α2β1-mediated cell migration [5], whereas both integrin α2β1 and DDR1 signaling are required for collagen-induced epithelial-mesenchymal transition in human pancreatic cancer BxPC-3 cells [6]. In addition, syndecan-1 supports integrin α2β1-mediated adhesion to collagen in human breast cancer MDA-MB-231 cells [7].

Collagen interacts with multiple biomacromolecules to mediate biological functions. Moreover, fibrillar collagens, exemplified by type I collagen, assemble into insoluble fibrils under physiological conditions. Given these properties, collagen function is typically investigated by assessing cellular responses to native collagen substrates while modulating receptor expression or activity. Such approaches include knockdown or overexpression of specific collagen-binding receptors, expression of mutant receptors, and the use of inhibitory antibodies. However, because simultaneous control of multiple collagen receptors remains technically challenging, current strategies are largely restricted to assessing the requirements or roles of individual receptors or crosstalk between selected receptor pairs.

The molecular basis of collagen interactions with collagen-binding biomolecules has primarily been elucidated using synthetic peptides that mimic the collagen triple-helical structure. For example, functional amino acid sequences within the triple helix that mediate interactions with collagen-binding molecules have been identified mainly using the Collagen Toolkit, a synthetic peptide library designed to systematically represent collagen segments [8]. Integrins α1β1, α2β1, α10β1, and α11β1 bind to the GxxGEx (x denotes any amino acid) sequence presented within the triple helix [9], whereas DDRs recognize the GVMGFO sequence [10,11] and HSPGs bind to the KGHRGF sequence [12]. Matricellular proteins also interact with collagen. Secreted protein acidic and rich in cysteine (SPARC) binds to the GVMGFO sequence [13,14] and von Willebrand factor (VWF) A3 domain interacts with the RGQOGVMGFO sequence [15], which overlaps with the DDR-binding sequence. Furthermore, collagen VII VWF A-like domain 2 recognizes the MGF sequence within the collagen triple helix, and this interaction has been proposed to be relatively weak and transient *in vivo* [16]. Pigment epithelium-derived factor (PEDF) interacts with the KGxRGFxGL sequence and shares binding sites with HSPGs in collagen [17].

Recently, integrins and DDRs were shown to cooperate in promoting osteoblast differentiation of the mouse calvaria-derived pre-osteoblast cell line MC3T3-E1 by culturing cells on plates coated with collagen-like triple-helical peptides containing either an integrin-binding or a DDR-binding sequence, alone or in combination [18]. These results highlight the utility of chemically synthesized collagen-like peptides as alternatives to native collagen. Unlike native collagen, which presents multiple signals at once, these peptides can selectively present defined signals to cells, enabling direct evaluation of their contributions to biological processes.

We previously developed peptide polymers (artificial collagens) composed of collagen-like triple-helical peptides [19]. These polymers are generated by end-to-end disulfide crosslinking of chemically synthesized triple-helical peptides. Building on these previous studies, we reasoned that copolymerizing multiple triple-helical peptides bearing distinct functional sequences in defined combinations and ratios would generate artificial collagen matrices with precisely tunable ligand composition. Using these substrates would enable systematic assessment of how the relative abundance of individual receptor-binding motifs influences cellular responses.

In this study, we used the PC12 cell line, derived from a rat adrenal medullary pheochromocytoma, as a model of neuronal differentiation to test this hypothesis. Upon exposure to nerve growth factor (NGF), PC12 cells differentiate into sympathetic neuron-like cells and extend neurites [20,21]. NGF induces differentiation *via* its receptor TrkA, and this process is enhanced when cells are cultured on collagen substrates [22]. Integrin α1β1 is required for PC12 cell differentiation on type I and type IV collagens [23]. Here, we used collagen-like peptide polymers to examine how collagen receptor-mediated inputs regulate NGF-dependent neuronal differentiation of PC12 cells. Our findings suggest opposing influences of collagen-binding receptors on this process, with integrin engagement promoting the differentiation response and DDR engagement suppressing it.

## 2. Materials and Methods

### 2.1. Peptide synthesis

Peptides were synthesized by 9-fluorenylmethoxycarbonyl-based solid-phase peptide synthesis on 2-chlorotrityl chloride resin (Peptide Institute, Osaka, Japan). Peptide resins were treated with a cocktail of trifluoroacetic acid (TFA)/*m*-cresol/thioanisole/3,6-dioxa-1,8-octanedithiol/H_2_O (80:5:5:5:5, v/v) for 4 h at room temperature. The desired peptides were purified using reversed-phase high-performance liquid chromatography (RP-HPLC) on a COSMOSIL 5C_18_-AR-II column (20 mm i.d. × 250 mm, Nacalai Tesque, Kyoto, Japan) with a linear gradient of acetonitrile in water, both containing 0.05% (v/v) TFA. The purified peptides were analyzed using RP-HPLC on a COSMOSIL 5C_18_-AR-II column (4.6 mm i.d. × 250 mm, Nacalai Tesque) (Fig. S1) and identified by electrospray ionization time-of-flight mass spectrometry (ESI-TOF MS; Compact, Bruker, Billerica, MA, USA) (Fig. S2). The synthesis and characterization of C3l-GPO, C3l-GFOGER, C3l-GVXGFO, and C3l-KGHRGF were previously reported [24].

### 2.2. Peptide polymerization

Synthesized peptides were dissolved in 0.05% (v/v) TFA in water at a final concentration of 1.11 mg/mL. The solutions were heated at 95 °C for 5 min, cooled at room temperature for 10 min, and incubated at 4 °C overnight to allow triple helix formation. The peptide solutions were mixed at the indicated ratios to prepare composite peptide solutions. Subsequently, dimethyl sulfoxide was added to each solution to a final concentration of 10% (v/v), and the mixtures were incubated at room temperature for 4 days to allow polymerization.

### 2.3. Cell culture

Rat pheochromocytoma PC12 cells were obtained from ATCC (CRL-1721). Cells were cultured in a 1:1 mixture of Dulbecco’s modified Eagle’s medium and Ham’s F-12 (DMEM/Ham’s F-12; FUJIFILM Wako Pure Chemical Corporation, Osaka, Japan) supplemented with 10% fetal bovine serum (FBS; Thermo Fisher Scientific, Waltham, MA, USA), 100 U/mL penicillin, and 100 µg/mL streptomycin (Sigma-Aldrich, St. Louis, MO, USA). Cells were maintained at 37 °C in a 5% CO_2_ atmosphere.

### 2.4. PC12 differentiation assay

Prepared peptide polymers and a type I collagen solution (Native Collagen Acidic Solution, IAC-30; KOKEN, Tokyo, Japan) were diluted to 50 µg/mL in sterile water. The 96-well plates (161093; Nunc, Thermo Fisher Scientific) were coated with 50 µL of each solution (2.5 µg/well) and dried overnight at room temperature. PC12 cells were pre-cultured in serum-free DMEM/Ham’s F-12 for 4 h prior to the assay. Cells were detached using 2 mM ethylenediaminetetraacetic acid (EDTA; FUJIFILM Wako Pure Chemical Corporation) in PBS and collected by centrifugation (120 × *g*, 24 °C, 4 min). Cells were resuspended to 4 × 10^4^ cells/mL in serum-free medium supplemented with 30 ng/mL NGF (Native mouse NGF 2.5S, N-100; Alomone Labs, Jerusalem, Israel) and 1 mM dibutyryl cyclic adenosine monophosphate (Bt_2_cAMP; Santa Cruz Biotechnology, Dallas, TX, USA). Each well was washed twice with PBS, and 100 µL of the cell suspension (4 × 10³ cells/well) was seeded into each well. Plates were incubated for 20 h at 37 °C in a 5% CO₂ atmosphere. Images were acquired from the central field of each well using a phase-contrast microscope (ECLIPSE TS100; Nikon, Tokyo, Japan). Differentiated cells were defined as those bearing at least one neurite with a length of ≥ 20 µm. The percentage of differentiated cells was calculated as the number of differentiated cells divided by the total number of cells per field. Images were analyzed using ImageJ software (version 1.54f; NIH, Bethesda, MD, USA).

### 2.5. Immunoprecipitation and western blotting

Six-well plates (140675; Thermo Fisher Scientific) were coated with 1.5 mL of 50 µg/mL solution of the prepared peptide polymers or type I collagen (75 µg/well) and dried overnight at room temperature. PC12 cells were pre-cultured and collected as described in Section 2.4. The cells were resuspended at 1.5 × 10^6^ cells/mL in serum-free medium supplemented with 30 ng/mL NGF and 1 mM Bt_2_cAMP.

Each well of the coated six-well plate was washed twice with PBS. The cell suspension (1.5 × 10^6^ cells/well) was seeded into each well and incubated for 4 h at 37 °C in a 5% CO₂ atmosphere. The cells were lysed with radioimmunoprecipitation assay (RIPA) buffer [1.0% (v/v) NP-40, 0.25% (w/v) sodium deoxycholate, 150 mM NaCl, 1 mM EDTA, and 50 mM Tris-HCl (pH 7.4)] supplemented with inhibitors (5 mM NaF, 1 mM sodium orthovanadate, 1 µg/mL pepstatin A, 1 µg/mL leupeptin, and 1 mM phenylmethylsulfonyl fluoride). The lysates were incubated on ice for 20 min and centrifuged (20600 × *g*, 4 °C, 15 min). Protein concentrations in the supernatants were determined using the Pierce BCA Protein Assay Kit (23227; Thermo Fisher Scientific).

For immunoprecipitation, cell lysates were incubated with anti-DDR1 rabbit monoclonal antibody (D1G6, 5583; Cell Signaling Technology, Danvers, MA, USA) or anti-DDR2 rabbit monoclonal antibody (E5S1S, 71991; Cell Signaling Technology) (1:150 dilution) at 4 °C for 14 h with rotation. Antigen-antibody complexes were captured with nProtein A Sepharose 4 Fast Flow (Cytiva, Marlborough, MA, USA) at 4 °C for 2 h with rotation. The beads were washed three times with RIPA buffer containing the inhibitors and collected by centrifugation (500 × *g*, 4 °C, 3 min). Immunoprecipitates were resuspended in 2 × SDS sample buffer [100 mM Tris-HCl (pH 6.7), 2% (w/v) SDS, 10% (v/v) glycerol, and 0.001% (w/v) bromophenol blue] containing 0.1 M dithiothreitol, and heated at 95 °C for 5 min. Immunoprecipitates prepared from equal amounts of lysate protein were subjected to SDS-PAGE using 8% polyacrylamide gels. Proteins were transferred to nitrocellulose membranes (Cytiva) by semi-dry transfer. The membranes were blocked with 5% skim milk, or 5% bovine serum albumin (BSA) in Tris-buffered saline [TBS; 50 mM Tris-HCl (pH 7.4) and 150 mM NaCl] at room temperature for 1 h. After two washes with TBS, the membranes were incubated with either the anti-DDR1 or anti-DDR2 antibody (1:400 dilution) in TBS containing 2% skim milk or anti-phosphotyrosine mouse monoclonal antibody (4G10, 96215S; Cell Signaling Technology; 1:400 dilution) in TBS containing 2% BSA at room temperature for 1 h. Membranes were washed three times with TBS and then incubated with horseradish peroxidase (HRP)-conjugated anti-rabbit IgG (goat, 7074P2; Cell Signaling Technology; 1:800 dilution) or HRP-conjugated anti-mouse IgG (goat, W402B; Promega, Madison, WI, USA; 1:1500 dilution) at room temperature for 30 min. Membranes were washed with TBS containing 0.1% Tween-20, developed using Pierce ECL Plus Western Blotting Substrate (Thermo Fisher Scientific) and imaged with a CCD imager LAS-3000 (FUJIFILM Corporation, Tokyo, Japan).

### 2.6. Statistical analysis

Results are presented as the mean ± SD, and individual data points are shown in the plots. Differentiation rates of PC12 cells were analyzed by one-way ANOVA followed by Tukey’s multiple-comparison test using GraphPad Prism version 7.04 (GraphPad Software, La Jolla, CA, USA).

## 3. Results

### 3.1. Design of the composition-controlled artificial collagens

To generate substrates with defined combinations and ratios of receptor-binding motifs, we used an artificial collagen platform based on end-to-end disulfide crosslinking of chemically synthesized triple-helical peptides (Fig. 1A). Each peptide chain comprised a repeating Gly–Xaa– Yaa sequence and contained three cysteine residues at each terminus. Functional sequences were introduced into a guest domain flanked by host domains consisting of repeated Pro–Hyp–Gly triplets that enhance the thermal stability of the triple helix. The peptide chains assembled into triple-helical structures in solution and were subsequently polymerized by disulfide crosslinking *via* the terminal cysteine residues.

**Figure 1.**
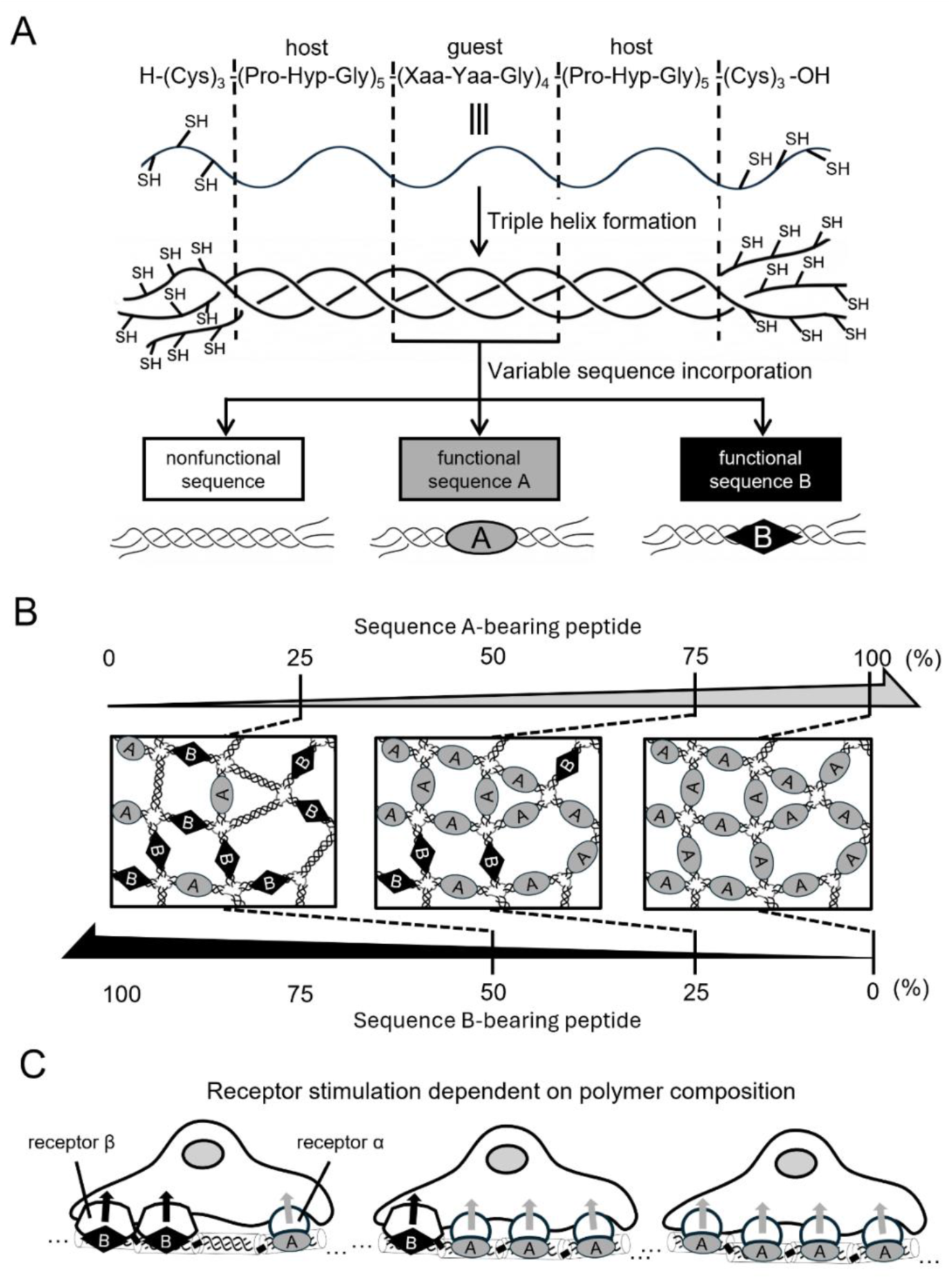
Schematic of the composition-controlled artificial collagen platform for analysis of collagen receptor crosstalk. (A) The peptide building blocks are chemically synthesized collagen-like peptides in which functional sequences (sequence A or sequence B) or a nonfunctional sequence can be introduced into a guest region. Each peptide comprises cysteine clusters at both termini flanking a triple helix-forming sequence and assembles into a triple-helical structure in solution. (B) Peptide polymers are generated by mixing triple-helical peptides bearing different sequences at defined ratios, followed by crosslinking *via* disulfide bond formation. By varying the proportions of peptides bearing sequence A and sequence B and supplementing the remaining fraction with the nonfunctional peptide, substrates with precisely controlled compositions of functional sequences are obtained. (C) Sequence A and sequence B mediate binding to receptor α and receptor β, respectively. Controlling the sequence composition of the peptide polymer enables assessment of individual receptor contributions and functional crosstalk during simultaneous receptor engagement.

Triple-helical peptides bearing distinct functional sequences were mixed at predetermined ratios before crosslinking and then polymerized under identical conditions (Fig. 1B). Importantly, the total peptide concentration was kept constant by supplementing the remaining fraction with a nonfunctional peptide (C3l-GPO) (Table 1). This design allowed the content of each functional motif to be independently tuned while maintaining a constant peptide-polymer background. The resulting composition-controlled peptide polymers provide a practical framework for assessing how multiple collagen receptor-mediated inputs are integrated to regulate cellular responses (Fig. 1C). We incorporated representative collagen-derived motifs that engage major classes of collagen-binding molecules: integrin-binding sequences (GLOGEN [25] and GFOGER [26,27]), an HSPG-binding sequence (KGHRGF) [17], and a DDR-binding sequence [GVXGFO; X = norleucine (Nle)]. In GVXGFO, the methionine residue of the native collagen motif GVMGFO was replaced with Nle, an oxidation-resistant analog [28,29]. The corresponding peptides were designated C3l-GLOGEN, C3l-GFOGER, C3l-KGHRGF, and C3l-GVXGFO, respectively (Table 1). The triple-helical conformation of each peptide at 37 °C was confirmed by the characteristic positive ellipticity at 225 nm in circular dichroism (CD) spectra (Fig. S3) [24,30].

**Table 1.** List of synthesized peptides.

| Peptide | Sequence |
| --- | --- |
| C3I-GPO | H-Cys-Cys-Cys-(Host)-Pro-Pro-Gly-Pro-Pro-Gly-Pro-Arg-Gly-Pro-Pro-Gly-(Host)-Cys-Cys-Cys-OH |
| C3I-GLOGEN | H-Cys-Cys-Cys-(Host)-Pro-Pro-Gly-Leu-Hyp-Gly-Glu-Asn-Gly-Pro-Pro-Gly-(Host)-Cys-Cys-Cys-OH |
| C3I-GFOGER | H-Cys-Cys-Cys-(Host)-Pro-Pro-Gly-Phe-Hyp-Gly-Glu-Arg-Gly-Pro-Pro-Gly-(Host)-Cys-Cys-Cys-OH |
| C3I-GVXGFO | H-Cys-Cys-Cys-(Host)-Pro-Arg-Gly-Gln-Hyp-Gly-Val-Nle-Gly-Phe-Hyp-Gly-(Host)-Cys-Cys-Cys-OH |
| C3I-KGHRGF | H-Cys-Cys-Cys-(Host)-Pro-Lys-Gly-His-Arg-Gly-Phe-Ser-Gly-Leu-Hyp-Gly-(Host)-Cys-Cys-Cys-OH |
| C3I-GVAGFO | H-Cys-Cys-Cys-(Host)-Pro-Arg-Gly-Gln-Hyp-Gly-Val-Ala-Gly-Phe-Hyp-Gly-(Host)-Cys-Cys-Cys-OH |

Host: (Pro-Hyp-Gly)<sub>5</sub>, Nle (X): norleucine

### 3.2. Collagen-binding integrins induce NGF-dependent differentiation of PC12 cells into neuron-like cells

To assess whether individual functional sequences derived from native collagen can independently promote differentiation of PC12 cells into neuron-like cells, we cultured PC12 cells for 20 h in the presence of NGF and Bt2cAMP on collagen-like peptide polymers containing a single functional sequence or on type I collagen (positive control). Peptide polymers containing 0, 3, 10, 30, or 100% of a peptide bearing a functional sequence were prepared by copolymerizing C3l-GPO with the corresponding peptide. Neurite outgrowth was not observed on uncoated surfaces or on the GPO polymer lacking functional sequences (Fig. 2A and S5). In contrast, marked neurite outgrowth was observed on type I collagen and on the 100% GLOGEN and 100% GFOGER polymers. Little to no neurite outgrowth was observed on the 100% GVXGFO and 100% KGHRGF polymers. The percentage of differentiated PC12 cells remained negligible on polymers containing GVXGFO or KGHRGF, regardless of the proportion of each sequence (Fig. 2B, definition of differentiated cell: see Materials and Methods). In contrast, the percentage of differentiated cells on GLOGEN- or GFOGER-containing polymers increased as the proportion of the respective sequence increased.

**Figure 2.**
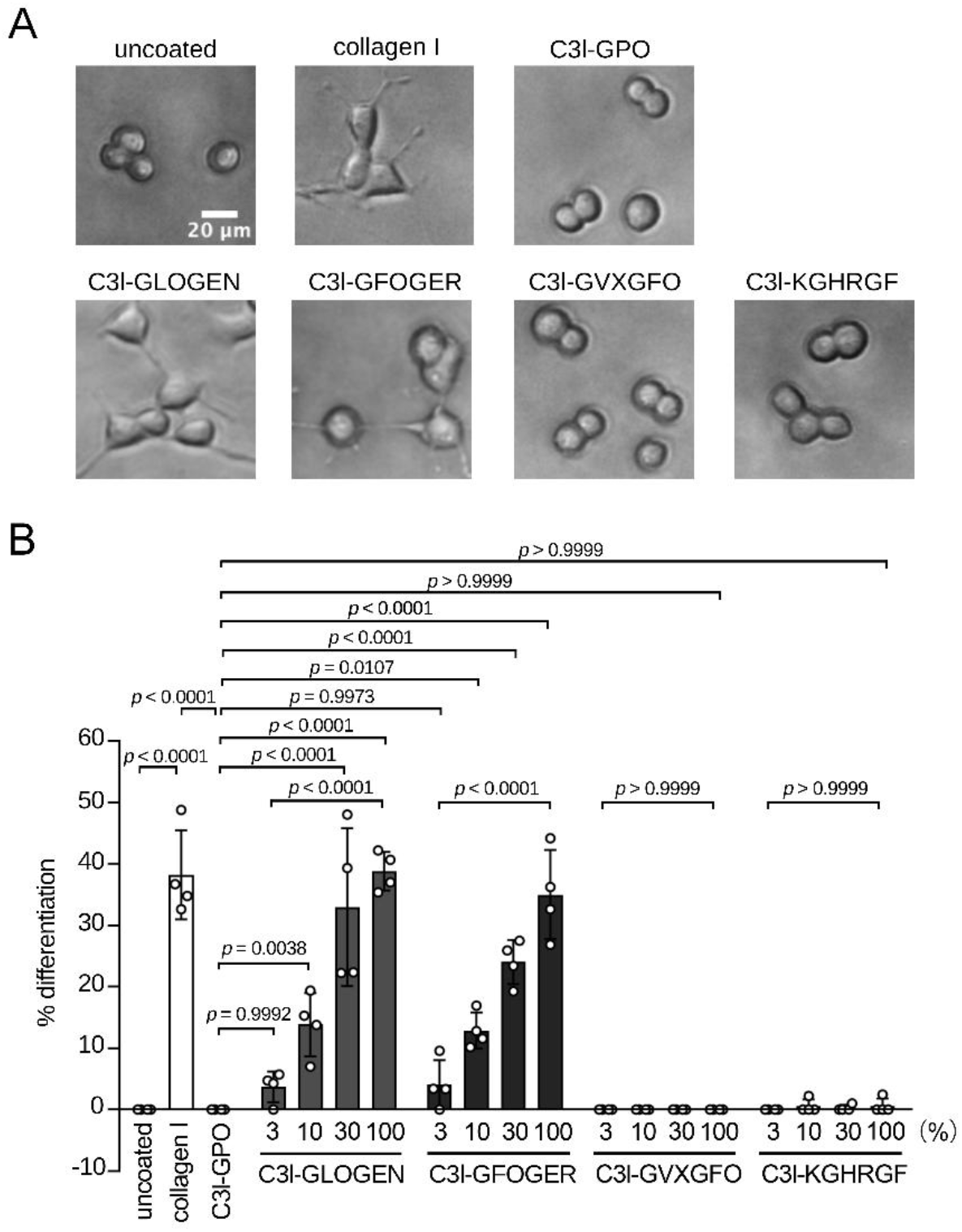
Peptide polymers containing integrin-binding sequences are sufficient to induce NGF-dependent differentiation of PC12 cells into neuron-like cells. (A) Representative phase-contrast images of PC12 cells cultured for 20 h in 96-well plates that were either uncoated or coated with type I collagen or the indicated single-component peptide polymers. Cells were treated with NGF and Bt^2^cAMP throughout the culture period. Images were captured using an inverted microscope. Brightness and contrast were adjusted uniformly across all images using ImageJ. (B) Percentage of differentiated PC12 cells after 20 h under the same conditions on the indicated substrates. Wells were coated with type I collagen, a nonfunctional peptide polymer (C3l-GPO), or peptide polymers containing the indicated proportions of functional sequence-bearing peptides. Data are presented as mean ± SD (n = 4), with each dot representing an individual well. P values were determined using one-way ANOVA followed by Tukey’s post-hoc test.

These findings indicate that signaling mediated by interactions between the collagen triple helix and collagen-binding integrins induces NGF-dependent differentiation of PC12 cells. Integrin α1β1 is the most likely integrin involved, as it has been reported to be the predominant collagen-binding integrin in PC12 cells and to be essential for their NGF-dependent neuronal differentiation [23,31,32].

### 3.3. The DDR-binding sequence suppresses integrin-induced NGF-dependent differentiation of PC12 cells into neuron-like cells

To examine whether other functional sequences modulate integrin-induced NGF-dependent differentiation of PC12 cells, we evaluated the differentiation of PC12 cells cultured on peptide copolymers containing GVXGFO or KGHRGF together with GLOGEN, which has a particularly high affinity for integrin α1β1 among collagen-binding integrins [25]. PC12 cells were cultured on peptide polymers consisting of 10% C3l-GLOGEN, 10–90% of either C3l-GVXGFO or C3l-KGHRGF, and the remainder C3l-GPO. The percentage of differentiated PC12 cells on polymers containing both GLOGEN and KGHRGF did not significantly differ from that on polymers containing GLOGEN alone (Fig. 3). In contrast, incorporation of GVXGFO into GLOGEN-containing polymers suppressed PC12 differentiation in a GVXGFO content-dependent manner. These findings suggest that signaling elicited by the GVXGFO sequence negatively regulates NGF-dependent neuronal differentiation mediated by integrin α1β1.

**Figure 3.**
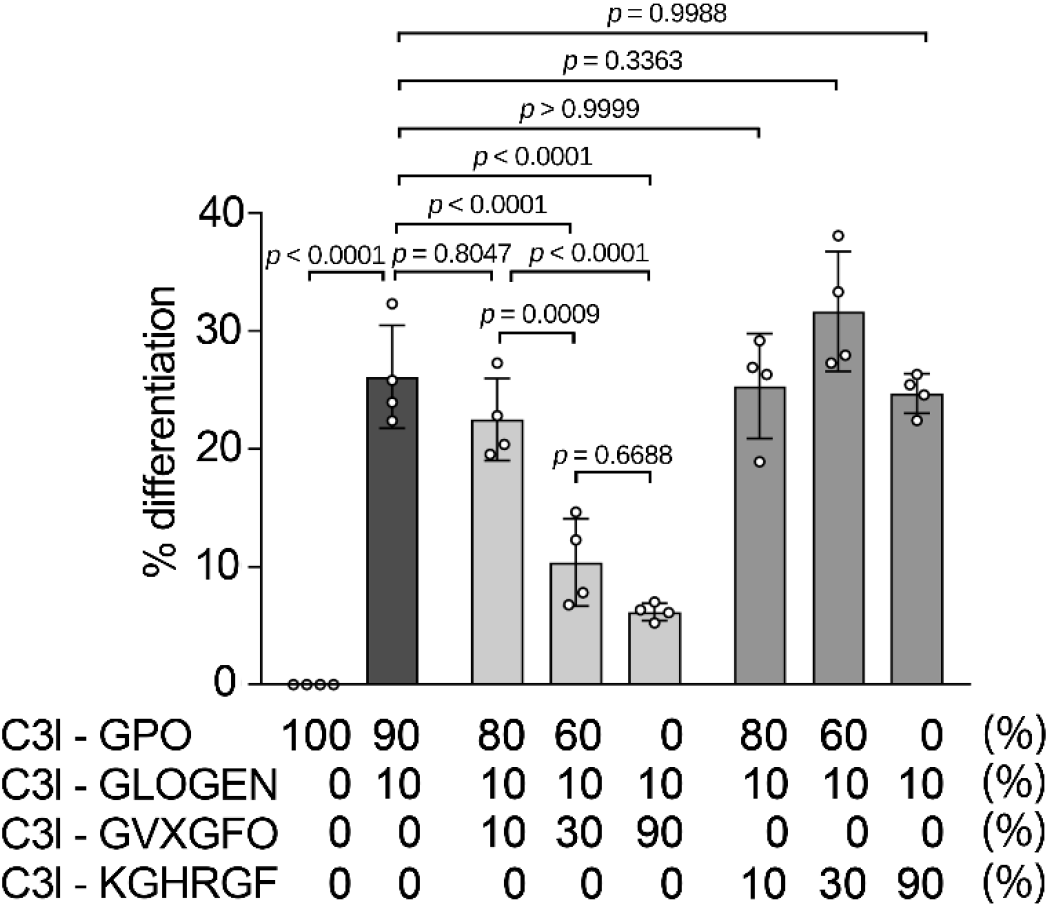
Integrin-induced differentiation of NGF-treated PC12 cells is suppressed by a DDR-binding sequence. Percentage of differentiated PC12 cells after 20 h in the presence of NGF and Bt₂cAMP on composite peptide polymers containing 10% C3l-GLOGEN together with 10, 30, or 90% C3l-GVXGFO or C3l-KGHRGF. Data are shown as mean ± SD (n = 4), with each dot representing an individual well. P values were determined using one-way ANOVA followed by Tukey’s post-hoc test.

### 3.4. Activation of DDRs is associated with the suppression of integrin-induced NGF-dependent neuronal differentiation of PC12 cells

To investigate the involvement of DDRs in the suppression of integrin-induced differentiation, we used composite peptide polymers containing C3l-GLOGEN together with either C3l-GVXGFO or C3l-GVAGFO (Table 1). Substitution of methionine with alanine in the GVMGFO sequence, yielding GVAGFO, has been reported to reduce its binding affinity for DDR1 and DDR2 [10,11], while maintaining its affinity for VWF [15]. GVXGFO inhibited GLOGEN-induced PC12 cell differentiation, whereas GVAGFO had no detectable effect (Fig. 4A). To explore candidate molecules that may contribute to the cellular responses to GLOGEN and GVXGFO, RT-PCR was performed to assess mRNA expression of cell-surface receptors and matricellular proteins capable of interacting with GLOGEN or GVXGFO in PC12 cells (Table S1 and Fig. S4). Among collagen-binding integrins known to bind GLOGEN, *Itga1* mRNA was detected under the culture conditions examined. Among the GVXGFO-binding proteins, *Ddr1, Ddr2,* and *Sparc* mRNAs were detected, whereas *Vwf* mRNA was not detected. These results suggest that DDRs and/or SPARC, but not VWF, may be involved in the suppression of PC12 cell differentiation.

**Figure 4.**
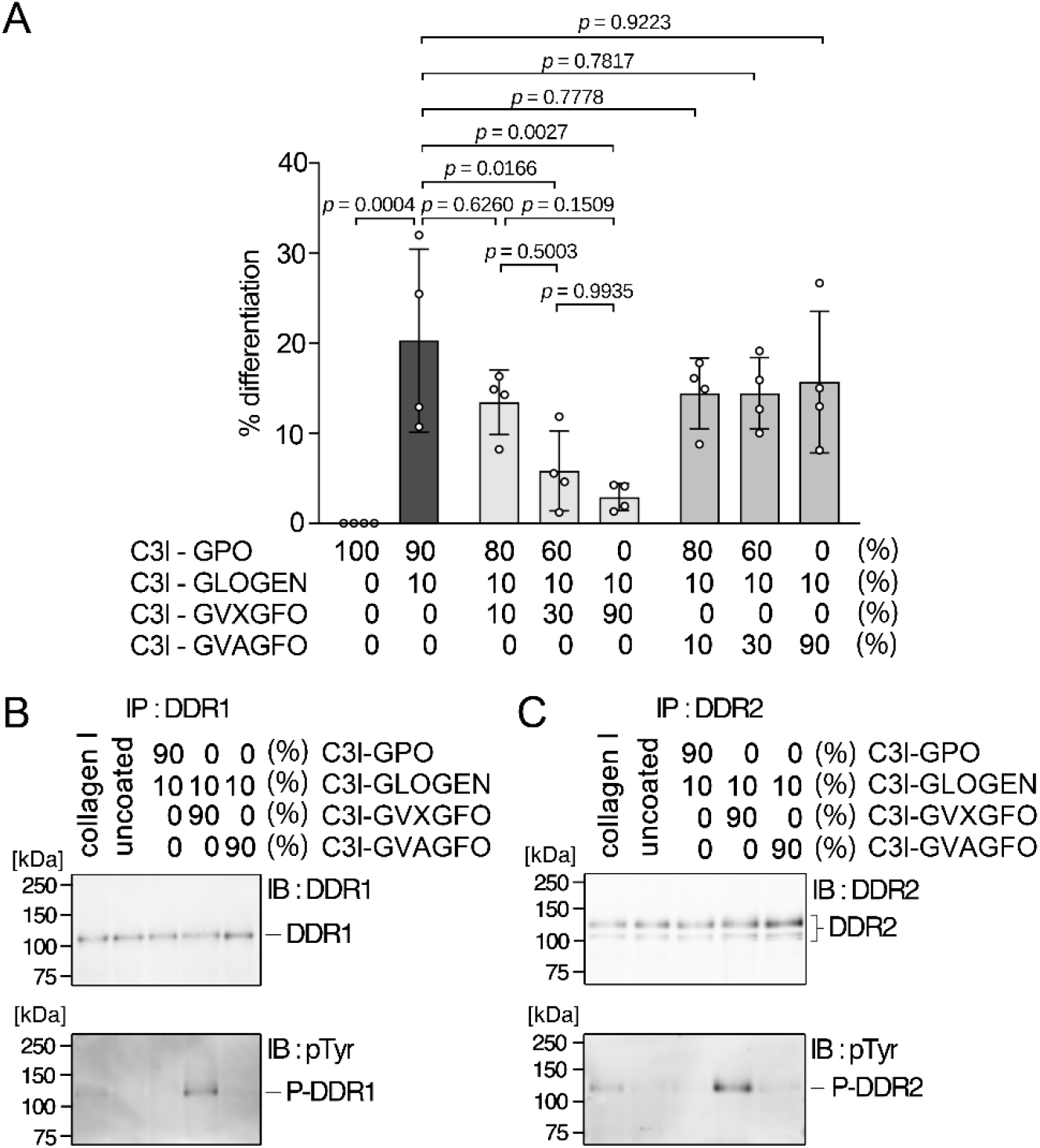
Suppression of integrin-induced differentiation of NGF-treated PC12 cells by a DDR-binding sequence is associated with DDR activation. (A) Percentage of differentiated PC12 cells after 20 h in the presence of NGF and Bt₂cAMP on composite peptide polymers containing 10% C3l-GLOGEN together with 10, 30, or 90% C3l-GVXGFO or C3l-GVAGFO. Data are presented as mean ± SD (n = 4), with each dot representing an individual well. P values were determined using one-way ANOVA followed by Tukey’s post-hoc test. (B, C) Immunoblot (IB) analysis of DDR1 (B) and DDR2 (C) tyrosine phosphorylation in PC12 cells cultured for 4 h in the presence of NGF and Bt₂cAMP on composite peptide polymers containing 10% C3l-GLOGEN together with 90% C3l-GPO, 90% C3l-GVXGFO, or 90% C3l-GVAGFO. DDR1 and DDR2 were separately immunoprecipitated (IP) from cell lysates and analyzed by immunoblotting for the corresponding DDR and phosphotyrosine (pTyr). P-DDR1 and P-DDR2 indicate tyrosine-phosphorylated DDR1 and DDR2, respectively.

Upon binding to collagen, DDRs undergo autophosphorylation at specific tyrosine residues, thereby activating downstream signaling pathways [33]. To determine whether DDR1 and DDR2 are activated, PC12 cells were cultured for 4 h in the presence of NGF on peptide polymers consisting of 10% C3l-GLOGEN and 90% C3l-GPO, C3l-GVXGFO, or C3l-GVAGFO. DDR1 and DDR2 were immunoprecipitated separately from cell lysates, and the precipitates were immunoblotted for phosphotyrosine (pTyr) and the corresponding DDR. DDR1 was detected in all DDR1 immunoprecipitates. DDR1 phosphorylation was observed in cells cultured on type I collagen and C3l-GLOGEN + C3l-GVXGFO polymers, but not in cells cultured on C3l-GLOGEN + C3l-GPO or C3l-GLOGEN + C3l-GVAGFO polymers (Fig. 4B). DDR1 phosphorylation was higher in cells cultured on C3l-GLOGEN + C3l-GVXGFO polymers than in those cultured on type I collagen. Similarly, DDR2 was detected in all DDR2 immunoprecipitates, whereas phosphorylated DDR2 was detected only in cells cultured on type I collagen and C3l-GLOGEN + C3l-GVXGFO polymers, with a stronger signal in the latter (Fig. 4C).

These results suggest that signaling through the cell-surface receptors DDR1 and/or DDR2 contributes to the suppression of integrin-induced NGF-dependent neuronal differentiation of PC12 cells.

## 4. Discussion

It remains challenging to disentangle the individual signals transmitted to cells through interactions between collagen and various biomolecules and to systematically assess the contribution of each signal to specific biological phenomena. In this study, we investigated receptor-specific biological activities of collagen and potential crosstalk among collagen receptors using artificial collagen polymers. These polymers were generated by crosslinking chemically synthesized triple-helical peptides bearing functional sequences in precisely controlled combinations and ratios.

A previous study reported that integrin α1β1 is required for the NGF-dependent differentiation of PC12 cells on collagen [23]. Here, we show that engagement of collagen-binding integrins alone is sufficient to induce neuronal differentiation of PC12 cells in the presence of NGF. Moreover, when an additional ligand sequence for another collagen-binding receptor was incorporated together with the integrin-binding sequence, the integrin-induced differentiation of PC12 cells was suppressed only in the presence of a sequence that binds to DDRs, and this suppression was associated with DDR phosphorylation. These findings suggest that neuronal differentiation may be regulated through crosstalk between DDRs and collagen-binding integrins. Notably, the relationship between integrins and DDRs appears to be context-dependent. While a previous study showed that integrins and DDRs cooperatively promote osteoblast differentiation of MC3T3-E1 pre-osteoblasts [18], our results suggest that DDR signaling opposes integrin-mediated neuronal differentiation of PC12 cells. Thus, the functional relationship between collagen receptors may vary among cell types and differentiation programs, allowing collagen to exert distinct effects on cell fate depending on cellular context.

To assess the potential contributions of specific integrin subtypes, we examined the expression of collagen-binding integrins in PC12 cells under our culture conditions. Consistent with previous reports, *Itga2* mRNA was not detectable in PC12 cells cultured without NGF [31], but became detectable upon NGF stimulation. We also detected *Itga10* and *Itga11* mRNAs in PC12 cells. Although collagen-induced PC12 differentiation has been reported to require integrin α1β1 [23], other collagen-binding integrin subtypes may also contribute to neuronal differentiation under our conditions. In addition, because these integrins recognize overlapping motifs in collagen [9], they may compete for binding to shared sites on the substrate, thereby modulating the relative contributions of integrin-dependent signals during neuronal differentiation.

DDRs are implicated in migration, proliferation, and survival, but their roles in the regulation of cell behavior remain incompletely defined. DDR1 is predominantly expressed in epithelial cells, whereas DDR2 is mainly expressed in stromal cells [2]. Both receptors recognize collagen types I, II, III, and V. Notably, DDR1 can also be activated by type IV collagen [34,35], whereas DDR2 recognizes type X collagen [36], indicating differences in ligand specificity and potentially in functional roles between the two receptors. In this study, it remains unclear whether DDR1, DDR2, or both receptors contribute to the suppression of NGF-dependent differentiation of PC12 cells. Further work will be required to clarify this point.

In this study, we used peptide polymers containing GVXGFO, in which the Met residue of GVMGFO is replaced with the isosteric non-natural amino acid Nle, as well as peptide polymers containing GVAGFO, in which the Met residue is substituted with Ala to suppress DDR binding [10,11]. These experiments demonstrate an important capability of artificial collagen matrices, in that diverse sequences can be incorporated irrespective of their abundance in, or even occurrence in, native collagen. Using this approach, we showed that a peptide polymer containing the GVXGFO sequence, previously reported to bind DDR2 with approximately 10-fold higher affinity than GVMGFO in solid-phase binding assays [28], can activate not only DDR2 but also DDR1 more effectively than native collagen. As additional receptor-selective motifs, including those for specific integrin subtypes [37], DDRs, and other collagen receptors, as well as higher-affinity sequences, are discovered or designed, artificial collagen may become an increasingly powerful tool for more precise control of cellular behavior.

The mechanism by which DDRs influence collagen-binding integrin function remains incompletely understood. DDRs are known to modulate integrin-mediated signaling through interactions with focal adhesion components [38,39]. Additionally, DDR- and integrin-mediated signals converge on multiple pathways, including the MAP kinase, PI3K/AKT, and Rho family GTPase pathways [40]. The MAPK/ERK1/2 pathway has been well characterized in PC12 cells and is activated upon NGF stimulation, playing a critical role in NGF-dependent neuronal differentiation [41]. Although some reports remain conflicting, the PI3K–Akt cascade is generally thought to promote neurite outgrowth in NGF-treated PC12 cells [42]. RhoA is a negative regulator of neurite outgrowth [43], whereas other Rho GTPases, such as Rac1 and Cdc42, are positive regulators [44]. A previous study also reported that multiple signaling pathways are involved in NGF-dependent differentiation of PC12 cells [45], and further investigation is required to elucidate the mechanism by which crosstalk between collagen-binding integrins and DDRs regulates neuronal differentiation. The artificial collagen-based approach described in this study offers a valuable means to investigate such collagen-mediated mechanisms.

Taken together, our findings demonstrate that composition-controlled artificial collagens provide a practical platform to probe how distinct collagen receptor inputs, alone and in combination, regulate cellular responses. This system should facilitate mechanistic studies of collagen-dependent cell regulation and support the design of tunable, cell-instructive biomaterials.

## 5. Conclusions

We prepared composition-controlled triple-helical peptide polymer substrates as artificial collagens by mixing collagen-like peptides bearing distinct receptor-binding sequences at predetermined ratios. The remaining fraction was supplemented with a nonfunctional peptide to maintain a constant total peptide content. In NGF-treated PC12 cells, integrin-binding sequences (GLOGEN or GFOGER) induced neuronal differentiation, whereas the DDR-binding sequence (GVXGFO) and the HSPG-binding sequence (KGHRGF) alone did not. Importantly, co-presentation of GVXGFO with an integrin-binding sequence suppressed neuronal differentiation relative to the integrin-binding sequence alone, and this suppression coincided with DDR1 and DDR2 phosphorylation. These findings have revealed opposing roles of collagen-binding integrins and DDRs in regulating NGF-dependent neuronal differentiation of PC12 cells, and established the artificial collagen system as a useful platform for dissecting how combinations of collagen receptor-mediated inputs regulate cellular responses.

## CRediT authorship contribution statement

**Kazunori K. Fujii:** Conceptualization, Methodology, Validation, Formal analysis, Investigation, Data Curation, Writing - Original Draft, Writing - Review & Editing, Visualization, Supervision, Project administration, Funding acquisition. **Kota Tsusaka:** Validation, Formal analysis, Investigation, Data Curation, Writing - Review & Editing, Visualization. **Takaki Koide:** Conceptualization, Methodology, Resources, Writing - Review & Editing, Supervision.

## Declaration of competing interest

The authors declare that they have no competing interests.

## Acknowledgments

This work was supported by a Waseda University Grant for Special Research Projects (Project No. 2022C-464) and JSPS KAKENHI Grant Number JP26K18467. We thank Yuito Hiura for assistance with experiments and data analysis. We also thank Rika Kawamura, Masahiro Nishikawa, Tamami Okajima, and Shinsuke S. Shibata for assistance with peptide synthesis. We further thank Shinichiro F. Ichise, Ryota Suzuki, and Hirotaka Tanaka for their support during the early stages of this work. We would like to thank Editage (www.editage.jp) for English language editing.

## Notes

### Competing Interest Statement

The authors have declared no competing interest.

