## Supplementary materials for "Composition-controlled artificial collagen shows opposing roles of collagen-binding integrins and discoidin domain receptors in neuronal differentiation of PC12 cells"

### Materials and Methods

#### Circular dichroism (CD) spectroscopy

CD spectra were recorded using a J-820 CD spectropolarimeter (JASCO, Tokyo, Japan) equipped with a Peltier temperature controller. The intensity of the spectra was converted to mean residue ellipticity ($[\theta]_{MRW}$). Peptides were dissolved at 0.5 mg/mL in 0.05% (v/v) trifluoroacetic acid (TFA) in water containing 10 mM tris(2-carboxyethyl)phosphine (TCEP). Samples were heated at 95 °C for 5 min, cooled at room temperature for 10 min, and incubated at 4 °C overnight. CD spectra were recorded from 190 to 260 nm using a quartz cell with a path length of 0.05 cm at 4, 37, and 85 °C.

#### RNA isolation and RT-PCR

PC12 cells were cultured in 35-mm dishes (150460; Thermo Fisher Scientific, Waltham, MA, USA), and total RNA was extracted using RNAzol RT (Molecular Research Center, Cincinnati, OH, USA) according to the manufacturer's instructions. The concentration and purity of the isolated RNA were assessed by measuring absorbance at 230, 260, and 280 nm using a NanoPhotometer N60 (Implen, Munich, Germany). For reverse transcription, 2 μg of RNA was used as a template in a reaction containing 0.5 mM each of deoxynucleotide triphosphate (dNTPs; Toyobo, Osaka, Japan) and 5 μM oligo$(dT)_{15}$ primer. The mixture was incubated at 65 °C for 5 min. Rat Universal Reference Total RNA (Zyagen, San Diego, CA, USA) was used as a positive control. After annealing, 200 U GeneAce Reverse Transcriptase (Nippon Gene, Tokyo, Japan) and 40 U RNase Inhibitor (Toyobo) were added on ice. Reverse transcription was performed in a total volume of 20 μL at 42 °C for 50 min, followed by heat inactivation at 70 °C for 15 min. Subsequently, 80 μL of Tris-EDTA [10 mM Tris-HCl (pH 8.0) and 1 mM EDTA] buffer was added to each sample. PCR was performed in a final volume of 20 μL containing 0.5 U rTaq DNA polymerase (Toyobo), 0.2 mM each dNTP, 0.5 μL of the cDNA synthesized above, and 1 μM of each gene-specific primer. Primer sequences for each gene are listed in Table S1. PCR was carried out under the following conditions. Initial denaturation was performed at 94 °C for 3 min. This was followed by 33 cycles of denaturation at 94 °C for 30 s, annealing at 65 °C for 30 s, and extension at 72 °C for 1 min. Final extension was performed at 72 °C for 10 min. PCR products were separated by electrophoresis on a 5% polyacrylamide gel in Tris-borate-EDTA buffer (TBE; 89 mM Tris, 89 mM boric acid, and 2 mM EDTA). Gels were stained with ethidium bromide and visualized under ultraviolet light at 365 nm.

**C3l-GLOGEN** $T_R$: 22.7 min **C3l-GVAGFO** $T_R$: 21.2 min

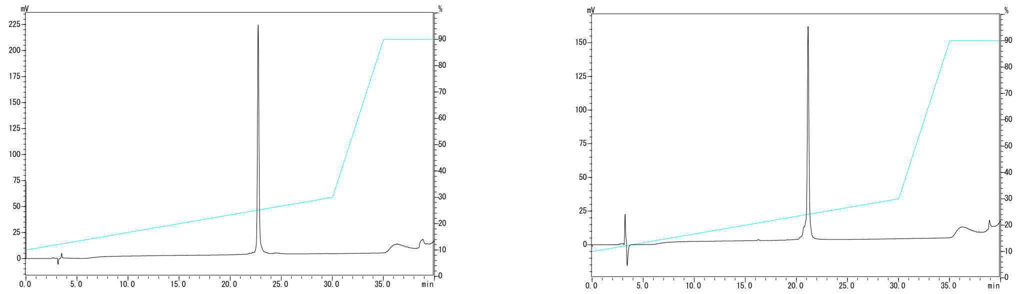

**Figure S1. High-performance liquid chromatography (HPLC) profiles of synthetic peptides.**

The peptides were analyzed using reversed-phase HPLC on a COSMOSIL $5C_{18}$-AR-II column (4.6 mm i.d. × 250 mm, Nacalai Tesque) at 60 °C. HPLC linear gradient: 10–30% acetonitrile in water, both containing 0.05% (v/v) TFA over 30 min, 30–90% over 5 min, and 90% for 5 min. Elution was monitored at 220 nm. The flow rate was 1 mL/min.

**C3l-GLOGEN**

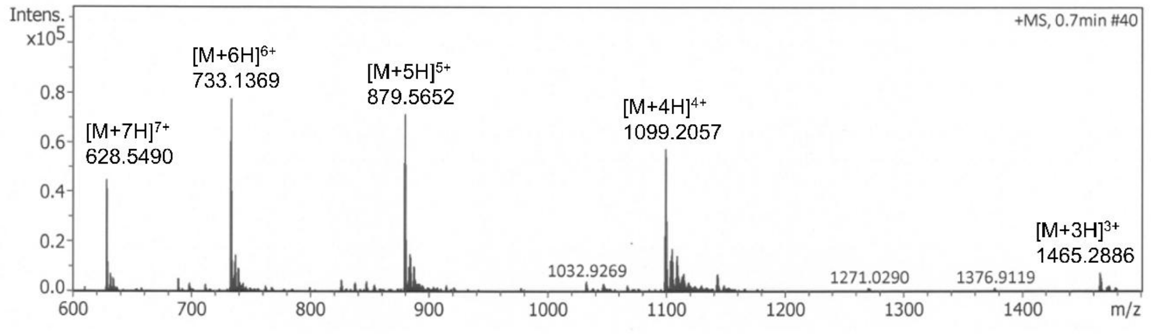


**C3l-GVAGFO**

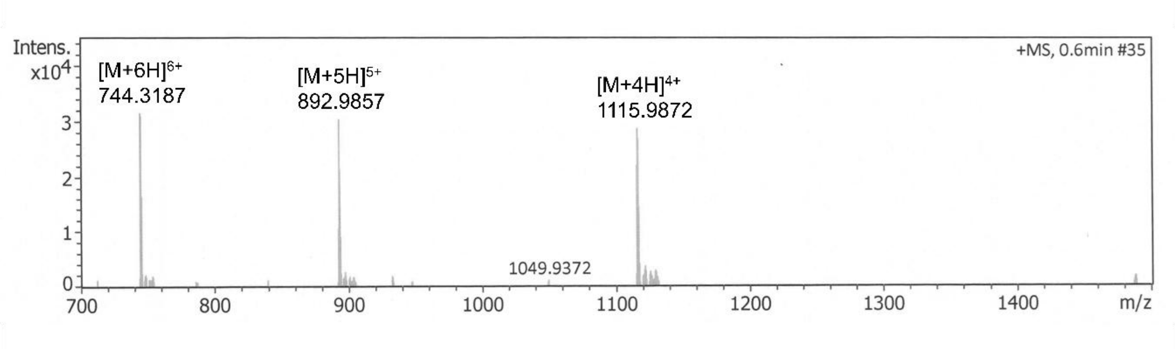


**Figure S2. Mass spectrometric analysis of the synthetic peptides.**

C3l-GLOGEN and C3l-GVAGFO were analyzed by ESI-TOF MS.

C3l-GLOGEN ESI-TOF MS (m/z): $[M + 7H]^{7+}$ calc. 628.5504 found 628.5490, $[M + 6H]^{6+}$ calc. 733.1409 found 733.1369, $[M + 5H]^{5+}$ calc. 879.5675 found 879.5652, $[M + 4H]^{4+}$ calc. 1099.2074 found 1099.2057, $[M + 3H]^{3+}$ calc. 1465.2739 found 1465.2886

C3l-GVAGFO ESI-TOF MS (m/z): $[M + 6H]^{6+}$ calc. 744.3164 found 744.3187, $[M + 5H]^{5+}$ calc. 892.9781 found 892.9857, $[M + 4H]^{4+}$ calc. 1115.9707 found 1115.9872

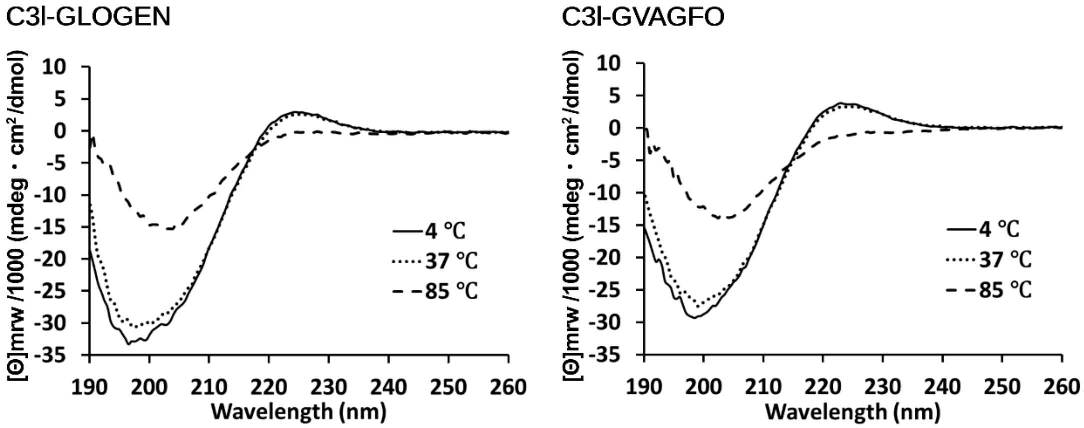


**Figure S3. CD analysis of the peptides.**

CD spectra were recorded on a J-820 CD spectropolarimeter equipped with a Peltier temperature controller at 4, 37, and 85 °C. Prior to measurement, peptides were annealed in 0.05% (v/v) TFA in water containing 10 mM TCEP.

**Table S1. Primers used for RT-PCR analysis**

| Gene | | Primer sequence (5′–3′) | Amplicon length [bp] | Ref |
|---|---|---|---|---|
| *Itga1* | Forward | GGCCCTGGTCACTATTGTTA | 184 | [1] |
| | Reverse | CATGACCACAGTTCCGTTCC | | |
| *Itga2* | Forward | CTAGCACTCCAACGGAGAGG | 200 | [2] |
| | Reverse | CACTGCACCTAGCATCAGGA | | |
| *Itga10* | Forward | GGCCTGTGCCCCTCTCTGGT | 123 | [3] |
| | Reverse | GGACAACGTTGGGCGGTCGG | | |
| *Itga11* | Forward | TGGAGGTCCAACACTTCCTC | 186 | [4] |
| | Reverse | GGGTTTCAGTCCCTCCTCTC | | |
| *Itgb1* | Forward | TTGGTCAGCAGCGCATATCT | 101 | [5] |
| | Reverse | ATTCCTCCAGCCAATCAGCG | | |
| *Ddr1* | Forward | ATGGAGCAACCATAGCTTCCC | 63 | [6] |
| | Reverse | CTCAGCCGGTCAAACTCAAACT | | |
| *Ddr2* | Forward | AGCGAGTCCAGCATGTTCAATAACA | 100 | [7] |
| | Reverse | GGTAGTCAGGGCGAAGGGGAA | | |
| *Vwf* | Forward | TCAAAGCCCCTGGACAACTC | 170 | [8] |
| | Reverse | TCCGAAAGGATTCATCTTGCC | | |
| *Sparc* | Forward | CCACTCGCTTCTTTGAGACC | 178 | [9] |
| | Reverse | TAGTGGAAGTGGGTGGGGAC | | |
| *Actb* | Forward | TCATGAAGTGTGACGTTGACATCCGT | 285 | [10] |
| | Reverse | CCTAGAAGCATTTGCGGTGCACGATG | | |

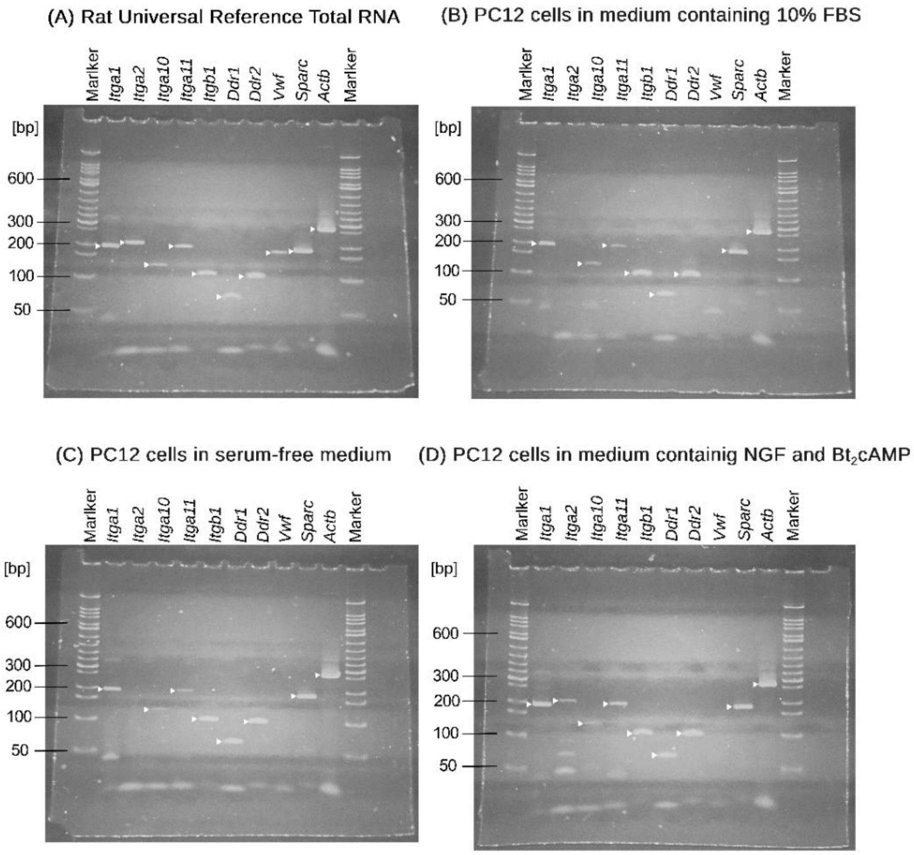


**Figure S4. Gel images of RT-PCR products for mRNAs encoding cell-surface receptors and extracellular proteins in PC12 cells cultured under various conditions.**

RT-PCR analysis of mRNA expression of cell-surface receptors and extracellular proteins capable of interacting with GLOGEN or GVXGFO. Each panel shows RT-PCR products obtained from total RNA isolated under the indicated conditions. Templates for reverse transcription were either (A) Rat Universal Reference Total RNA (positive control) or (B–D) total RNA isolated from PC12 cells cultured under the following conditions: (B) in medium containing 10% FBS; (C) for 4 h after replacement of the medium with serum-free medium; or (D) for 4 h in serum-free medium, after which the medium was replaced with medium containing 30 ng/mL NGF and 1 mM $Bt_2$cAMP, followed by a further 20 h of culture. Reverse transcribed cDNA was amplified by PCR, and the products were separated by electrophoresis on 5% polyacrylamide gels and visualized by ethidium bromide staining. White arrowheads indicate bands corresponding to the expected PCR product sizes. A 50-bp DNA ladder was used as a size marker.

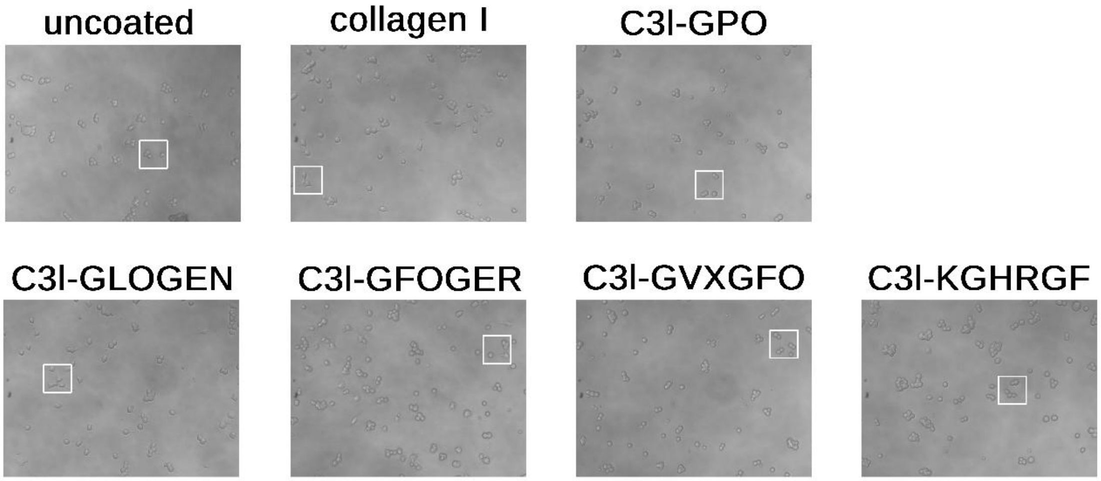


**Figure S5. As-acquired, uncropped phase-contrast images used for Fig. 2A.**

Images were acquired using an inverted microscope. The regions shown in Fig. 2A are indicated by white squares.
